# Dicamba drift alters patterns of chewing herbivory in three common agricultural weeds

**DOI:** 10.1101/2022.08.21.504705

**Authors:** Nia Johnson, Grace Zhang, Anah Soble, Regina S Baucom

## Abstract

How agricultural regimes, such as novel herbicide exposure, may influence plant-herbivore interactions and specifically patterns of plant herbivory has come under increased interest in recent years due to rapidly changing herbicide use in agroecosystems. This paper examines patterns of plant herbivory using three common agricultural weeds exposed to low doses of dicamba, a synthetic auxin herbicide that is exponentially increasing in use given the adoption of dicamba tolerant crops. We used a replicated field study to examine how the amount and type of chewing herbivory may be altered in *Ipomoea purpurea* (common morning glory, Convolvulaceae), *Datura stramonium* (jimsonweed, Solanaceae), and *Abutilon theophrasti* (velvetleaf, Malvaceae) exposed to dicamba drift (*i.e*., 1% of the field dose). We found an increase in chewing herbivory damage when plants were exposed to dicamba and changes in the type of herbivory following exposure. Chewing herbivory differed among species in the presence of dicamba drift: *A. theophrasti* and *D. stramonium* showed more total leaf-chewing herbivory than controls, but *I. purpurea* showed no difference in the overall amount of herbivory. We also found that the type of herbivory was significantly altered in drift. *A. theophrast*i and *I. purpurea* both exhibited declines in hole feeding but increases in margin feeding, whereas *D. stramonium* showed no such changes. Overall, our results show that herbicide drift can induce shifts in plant-herbivore interactions, highlighting the need for mechanistic studies to uncover the cause underlying the shifts and comparative studies on weed communities to understand long-term consequences.

## INTRODUCTION

The interactions between plants and their herbivores are central to both ecological and evolutionary dynamics. Herbivores consume between 5-20% of the plant biomass produced annually (Cyr and Face, 1993; Agrawal, 2011; Turcotte et al., 2014), making plant herbivory a significant effector of ecosystem dynamics, stability, productivity, and nutrient balance (Schmitz, 2008; Tilman et al., 2012), as well as an important driver of evolutionary processes (Futuyma and Agrawal, 2009). How plant-herbivore interactions may be altered as a consequence of novel abiotic stressors – especially those associated with scenarios of global change, urbanization, and land use for agriculture (Massad and Dyer, 2010; Turcotte et al., 2017; Miles et al., 2019) – has come under increased scrutiny given the central role plant herbivory plays on the ecological and evolutionary dynamics of communities (Huntly, 1991).

The expansion and intensification of agriculture has significantly altered Earth’s landscapes, with agriculture comprising approximately one-third of ice-free land area (Ramankutty et al., 2018). Within agricultural landscapes, herbicide use leads to a number of unintended consequences, such as the evolution of herbicide resistance (Baucom, 2019), dramatic changes to weed community structure (*i.e*., ‘weed shifts’; Owen, 2008), and the exposure of nontarget organisms to herbicide drift, typically defined as the unintentional exposure of nontarget areas to low doses of herbicide (∼1-5% of the field dose, Cessna et al., 2005). How herbicide drift may impact weed communities, the insects they support, and plant herbivory are each critical gaps in our knowledge given the high potential for herbicide drift in agroecosystems and the central role plant herbivory plays in ecological and evolutionary processes. Unfortunately, our understanding of the effect of unintentional herbicide exposure on plant-herbivore interactions remains limited.

For example, while there are a number of studies that assess how direct exposure to herbicides may impact the survival and growth of insects (Bohnenblust et al., 2013; Sharma et al., 2018; Fuchs et al., 2020), there are strikingly few reports of how herbicides impact insect herbivory (but see Gutiérrez et al., 2020), and few that examine plant herbivory in the context of herbicide drift (but see Johnson and Baucom 2022). This dearth of knowledge is especially relevant given the recent and widespread shift to the herbicide dicamba (3,6-dichloro-2-methoxybenzoic acid) in contemporary agriculture (Wechsler et al., 2019). Dicamba, a synthetic auxin herbicide, is increasing exponentially following the recent adoption of dicamba tolerant crops (*i.e*., 600% increase between 2014 and 2019 in the US alone; Baker, 2022). This herbicide is also known for drifting post-application (Egan et al., 2014a; Jones et al., 2019), and the number of unintentional drift events has likewise increased along with its use (EPA, 2021).

Because dicamba is a synthetic auxin, exposure at low doses may be expected to disrupt or alter patterns of plant herbivory. Auxin is a naturally produced plant phytohormone that is found throughout plant structures and the plant life cycle (Teale et al., 2006). Synthetic auxins act as herbicides by deregulating normal patterns of plant growth, ultimately causing death by distorting cell division and leading to a collapse of tissue structure (Grossmann, 2010). Exposure to synthetic auxins through an herbicide drift event may directly impact herbivory defense responses since natural auxins closely interact with other phytohormones involved in defense – jasmonic acid (JA), ethylene and abscisic acid (ABA) (Peña-Cortés et al., 1995; Liu and Wang, 2006; Muday et al., 2012; Xu et al., 2020). Despite the likely role of dicamba exposure on plant herbivory, our knowledge of how dicamba drift may affect patterns of herbivory in weed communities remains exceedingly limited: while one study shows an increased abundance of *Bemisia tabaci* (whitefly) larvae, and an (albeit nonsignificant) increase in chewing herbivory on plants exposed to dicamba drift (Johnson and Baucom, 2022), there are no other examples, to our knowledge, that examine the influence of dicamba drift on herbivory.

Here, we investigate the effect of dicamba drift on chewing herbivory in three common weeds of agriculture – *Ipomoea purpurea* (common morning glory), *Datura stramonium* (jimsonweed), and *Abutilon theophrasti* (velvetleaf). We elected to examine three species since herbivore dietary breadth and composition are specialized by individuals, populations, and species (Forister et al., 2015), making studies with multiple plant hosts particularly valuable for understanding complex interactions within these communities. Specifically, we performed a replicated field study to determine if the amount of herbivory damage was altered when plants were exposed to dicamba drift. Moreover, due to the diversity of insect herbivores that produce distinct patterns of consumption (Labandeira, et al., 2005; Carvalho et al., 2014), we also quantified the types of leaf-chewing damage to capture potential shifts in insect community structure. Our specific questions were: 1) Does dicamba drift alter the proportion or type of chewing herbivory experienced by plants, and 2) Do different weed species show different patterns of chewing herbivory when exposed to dicamba drift? Given that recent work in *A. theophrasti* showed a non-significant increase in chewing herbivory (Johnson and Baucom, 2022), we expected to find similar results for this species; however, we had no *a priori* expectation for how patterns of herbivory might be altered in the other species examined, especially since they are in separate families (Convolvulaceae, Solanaceae, and Malvaceae) and are thus each distantly related from one another.

## MATERIALS AND METHODS

### Field experiment

On July 14, 2018, we planted replicate seeds of three common weed species – *Ipomoea purpurea* (common morning glory), *Datura stramonium* (jimsonweed), *Abutilon theophrasti* (velvetleaf) – into two field plots separated by 50 m at the Matthaei Botanical Gardens in Ann Arbor, MI. Within each plot, we randomized two replicate seeds per maternal line per species within two treatments (dicamba drift and control) randomly assigned in two separate blocks. In total we planted 320 replicate seeds (2 plots x 2 blocks x 2 treatments x 2 replicates x 20 maternal lines (10 *I. purpurea*, 5 *D. stramonium*, 5 *A. theophrasti*)). Lines used in the experiment were originally sampled as seed from agricultural fields from either the southeast (*I. purpurea*) or midwest (*A. theophrasti* and *D. stramonium*) and stored as seed until used in the experiment. Although sampled from agricultural fields, meaning that lineages likely experienced some herbicide exposure, we do not have a history of specific herbicide exposure. However, the *I. purpurea* lines used in the study were sampled in 2012, and both of the other species were sampled in 2015, making it likely that none of the lineages used in this study were exposed to dicamba, since this herbicide has only recently increased in use given the release of dicamba tolerant crops in 2016 (Wechsler et al., 2019).

After five weeks of growth, we recorded the number of leaves of each individual for a pre-treatment size estimate and then applied 1% the field dose of dicamba (5.6 g ai/ha) to half of the experimental individuals. Plants were sprayed from one pass until just wet using a handheld multi-purpose sprayer set to a medium-fine mist with an operating pressure of 45 PSI. We elected to use a handheld multipurpose sprayer rather than a CO_2_ sprayer, which maintains a constant application rate, to simulate the variation that would occur in light of an off-target drift event in field conditions. Two weeks after treatment with dicamba we recorded the total number of leaves and the number of leaves that displayed evidence of dicamba damage (*i.e*., leaf cupping/malformation, a type of damage that looks very distinct).

At the end of September, we randomly sampled 4-5 leaves from a subsample of 126 randomly chosen plants from each block and treatment combination and scanned the images using a scanner (Canon CanoScan L110) to document both the amount and type of chewing herbivory plants experienced in the field. We used the imaging software Fiji ImageJ (vers 3.0; Schindelin et al., 2012) to obtain total leaf area and chewing damage. To do so, we converted each image to binary format and used the particle analysis tool to convert total pixel number into centimeters for a measure of leaf area, and then used a macro plug-in to place a grid with 0.1cm^2^ spacing over each image and counted number of grids with obvious chewing herbivory damage to estimate the amount of herbivore damage per leaf. Our measure of proportion herbivory is the total area in pixels with chewing herbivory damage divided by the total leaf area for each leaf, and for data analysis, we averaged the proportion of chewing herbivory per individual.

We recorded the presence of each type of chewing herbivory damage per leaf, first focusing on whether or not the damage was from hole, margin, or surface feeders, and then further refined the leaf damage chewing types into 18 distinct damage types (following the *Guide to Insect (and other) Damage Types on Compressed Plant Fossils* (Labandeira, et al., 2005). We statistically evaluated the potential for changes in the type of chewing herbivory damage across the three main categories (hole, margin, and surface) and present differences according to the refined categories for illustrative purposes. To analyze herbivory type changes, we developed a proportion of leaf herbivory type by assessing the number of leaves per plant that displayed any of the three types of damage and divided this by the total number of leaves categorized per individual (typically 4 leaves per plant). Although we did not quantify herbivore taxonomic groups in this study, some of the leaf-chewing groups observed on experimental plants were Coleoptera (beetles), Orthoptera (grasshoppers), Phasmida (stick insects), and Lepidoptera (caterpillars).

### Data analysis

We used R vers 4.2.0 (Team, 2016) in RStudio (RStudio Team, 2020) for all analyses and generated figures using ggplot 2 vers 3.3.5 (Wickham, 2016). We identified the most appropriate transformation for the proportion leaf herbivory using the bestNormalize package (Peterson et al., 2021) to meet the assumptions of ANOVA. Our main interests were to determine if the proportion of chewing herbivory present on leaves was affected by the presence of dicamba drift, and if the three species exhibited differences in the proportion of herbivory according to treatment. Thus, we used log transformed proportion herbivory as the dependent variable in a mixed model analysis of variance with species, treatment (dicamba or control), block, and the interaction between species and treatment as the independent variables using the lmer function of the lme4 package vers 1.1-29 (Bates et al., 2011).

All independent variables were considered fixed except for block, which was modeled as a random effect. Degrees of freedom were calculated using Satterthwaite’s method (Satterthwaite, 1946). We performed planned comparisons of the treatment effect within each species using a non-parametric Kruskal Wallis test. We next determined if the type of chewing herbivory was altered in the presence of drift by using log transformed proportion of leaves with each type of damage as a dependent variable and damage type (hole, margin, or surface), treatment (dicamba or control), block, and species and their interactions as independent variables in a mixed model analysis of variance, again using the lmer package. As above, all independent variables were considered fixed except for block, which was modeled as a random effect.

## RESULTS

Across plants, a low percentage of leaf material, on average, showed chewing leaf herbivory (*i.e*., 0-3% per leaf) in this experiment. However, we found a significant effect of dicamba drift on the average proportion of herbivory compared to control conditions (Avg ±se dicamba: 0.41% ±0.06 vs control: 0.11% ±0.02; Treatment effect, F-value = 20.13, p-value < 0.001; Table 1). There was no detectable difference in the average proportion of herbivory experienced by species (*i.e*. a non-significant species effect; Table 1) nor was there a significant species by treatment interaction (Table 1). However, within-species planned comparisons of the proportion herbivory showed that *A. theophrasti* and *D. stramonium* clearly experienced more herbivory in the dicamba treatment compared to the control environment (*A. theophrasti*: χ^2^_1_ = 17.62, p-value < 0.0001; *D. stramonium*: χ^2^_1_ = 5.10, p-value = 0.02; Fig 1) whereas *I. purpurea* did not (χ^2^_1_ = 0.25, p-value = 0.62; Fig 1). We additionally uncovered a significant effect of Block on the average proportion of herbivory (Table 1), meaning that there was unexplained environmental variation influencing the overall amount of herbivory experienced by plants.

**Table 1.** Mixed model analyses of variance examining the A) average proportion herbivory from a sample of 3-4 leaves per individual, and B) the proportion of leaves showing different types of herbivory, *i.e*., hole, margin, or surface damage.

| **A. Proportion herbivory** |  |  |  |  |  |
| --- | --- | --- | --- | --- | --- |
| *Fixed effects* | MeanSq | Numdf | Dendf | F-value | p-value |
| Species | 0.01 | 2 | 117.78 | 0.19 | 0.82 |
| Treatment | 1.02 | 1 | 117.13 | 20.13 | **<0.001** |
| Species × treatment | 0.08 | 2 | 118.04 | 1.59 | 0.21 |
| *Random effects* | | | | $\chi^2$ | |
| Block |  | 1 |  | 3.77 | **0.05** |
| **B. Proportion leaves** |  |  |  |  |  |
| *Fixed effects* | MeanSq | Numdf | Dendf | F-value | p-value |
| Herbivory type | 0.33 | 2 | 54 | 102.09 | **<0.001** |
| Species | 0.00 | 2 | 54 | 0.67 | 0.51 |
| Treatment | 0.00 | 1 | 54 | 0.39 | 0.54 |
| Herbivory type × species | 0.05 | 4 | 54 | 14.82 | **<0.001** |
| Herbivory type × treatment | 0.19 | 2 | 54 | 59.19 | **<0.001** |
| Species × treatment | 0.00 | 2 | 54 | 0.73 | 0.49 |
| Herbivory type × species × treatment | 0.04 | 4 | 54 | 12.45 | **<0.001** |
| *Random effects* | | | | $\chi^2$ | |
| Block |  | 1 |  | 0.00 | 1 |

**Fig 1.**
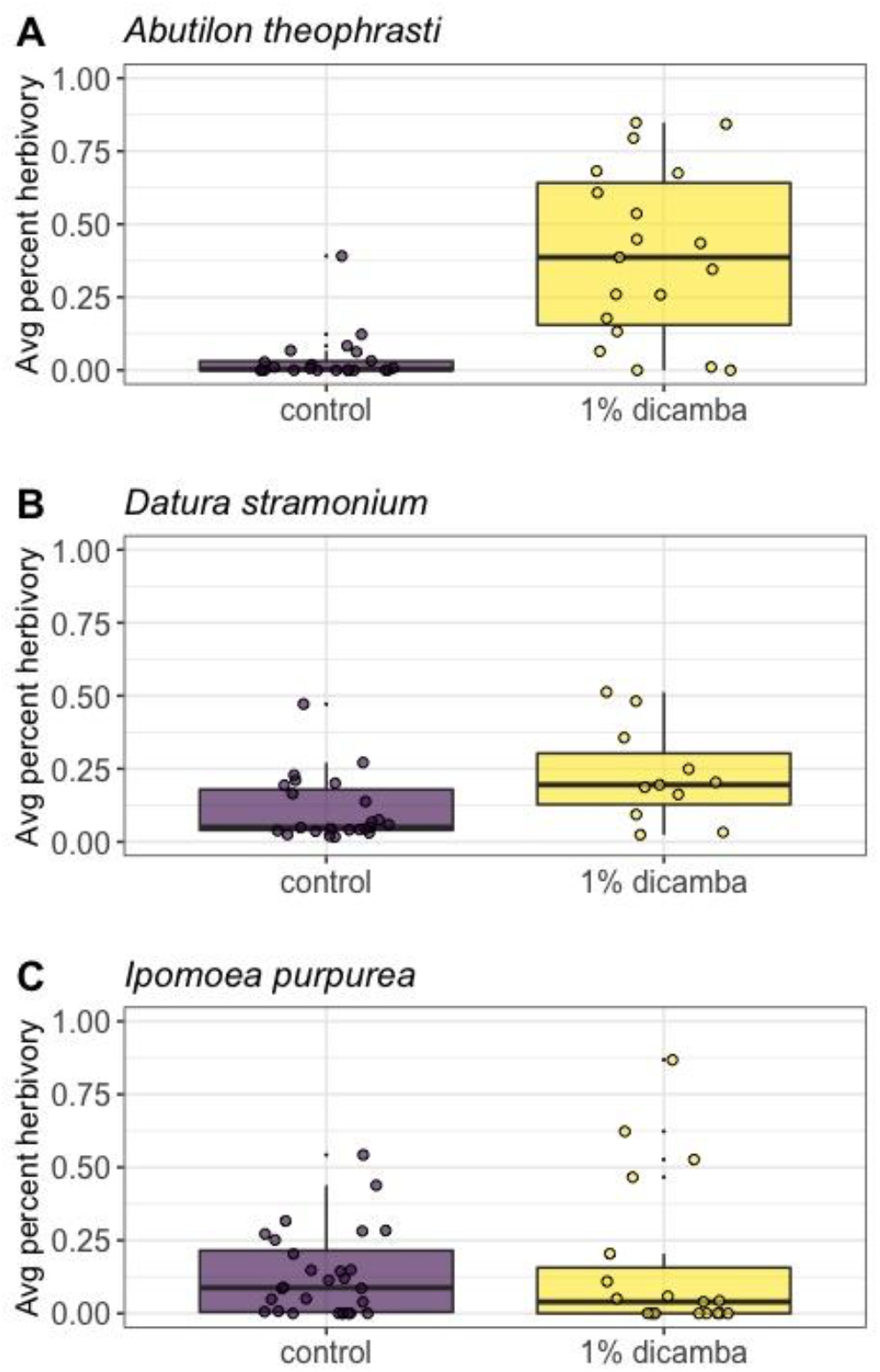
The average percent herbivory across replicate plants of A) *Abutilon theophrasti* (velvetleaf), B) *Datura stramonium* (jimsonweed), and C) *Ipomoea purpurea* (common morning glory) experienced in field conditions in the presence of 1% the field dose of the herbicide dicamba (5.6 g ai/ha) or in non-dicamba control environment. The proportion of each leaf showing herbivory was recorded on replicate leaves of each individual using ImageJ, averaged per plant, and shown here as a percentage.

We next examined the potential that the type of herbivory experienced by plants (either hole, margin, or surface feeding) differed according to exposure to dicamba drift. We found significant differences in the types of herbivory identified on leaves across species (Table 2), with a higher proportion of leaves showing hole and margin damage compared to surface feeding (Fig 2). While there was no evidence for an overall treatment effect (F-value = 0.39, p-value = 0.54), we found a highly significant interaction between herbivory type and species (F-value = 14.82, p-value < 0.001), indicating that the different species exhibited different types of herbivory. We also found a significant interaction between herbivory type and treatment (F-value = 59.19, p-value < 0.001), meaning that dicamba application altered the type of herbivory experienced overall, and finally, there was a significant three-way interaction between herbivory type, treatment, and species (F-value = 12.45, p-value < 0.001), indicating that the type of herbivory differed according to species and dicamba exposure. For both *A. theophrasti* and *I. purpurea*, the proportion of leaves that exhibited hole feeding was significantly lower in the dicamba treatment compared to the control environment (*A. theophrasti*, average±se : 0.13±0.07 dicamba vs 0.82±0.10 control; *I. purpurea*: 0.40±0.11 dicamba vs 0.79±0.08 control) whereas the proportion of leaves showing margin feeding increased substantially in the presence of dicamba for both species (*A. theophrasti*: 0.89±0.06 dicamba vs 0.14±0.08 control; *I. purpurea*: 0.73±0.09 dicamba vs 0.41±0.09 control; Fig 2). Furthermore, in *A. theophrasti* we also identified a significantly lower proportion of leaves showing surface feeding in the presence of dicamba compared to control conditions (0.03±0.03 dicamba vs 0.25±0.11 control; Fig 2) but did not find substantial changes in this type of feeding for the other two species. The other types of herbivore chewing damage – hole and margin – likewise did not differ between treatments for *D. stramonium* (Fig 2).

**Fig 2.**
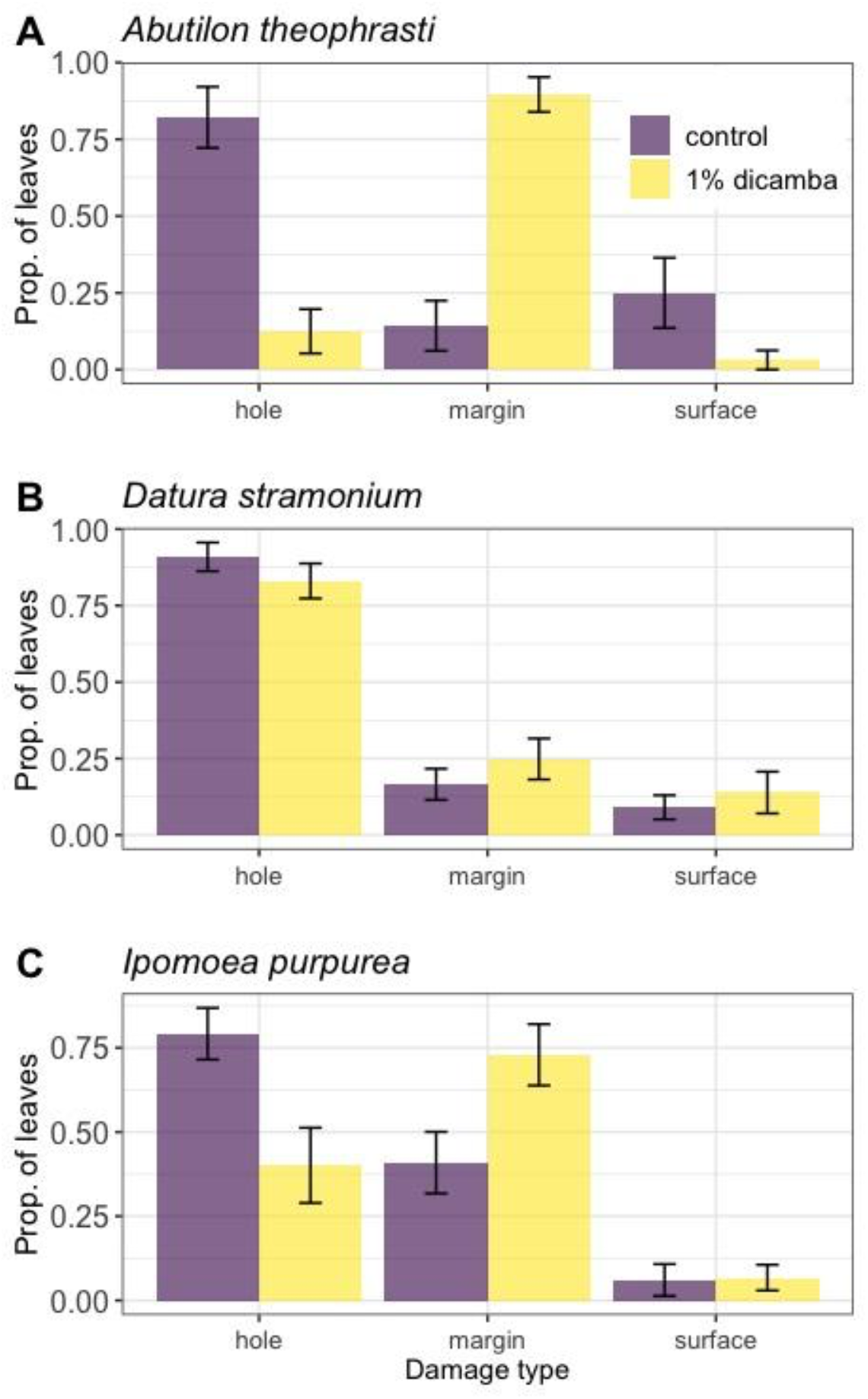
The proportion of leaves exhibiting either hole, margin, or surface herbivory across replicate plants of A) *Abutilon theophrasti* (velvetleaf), B) *Datura stramonium* (jimsonweed), and C) *Ipomoea purpurea* (common morning glory) growing in field conditions and treated with 1% the field dose of the herbicide dicamba (5.6 g ai/ha) or a non-dicamba control environment. The type of herbivory was recorded on three or four leaves of each species, and the average proportion of each herbivory type (± standard error) across plants of each species is shown.

**Fig 3.**
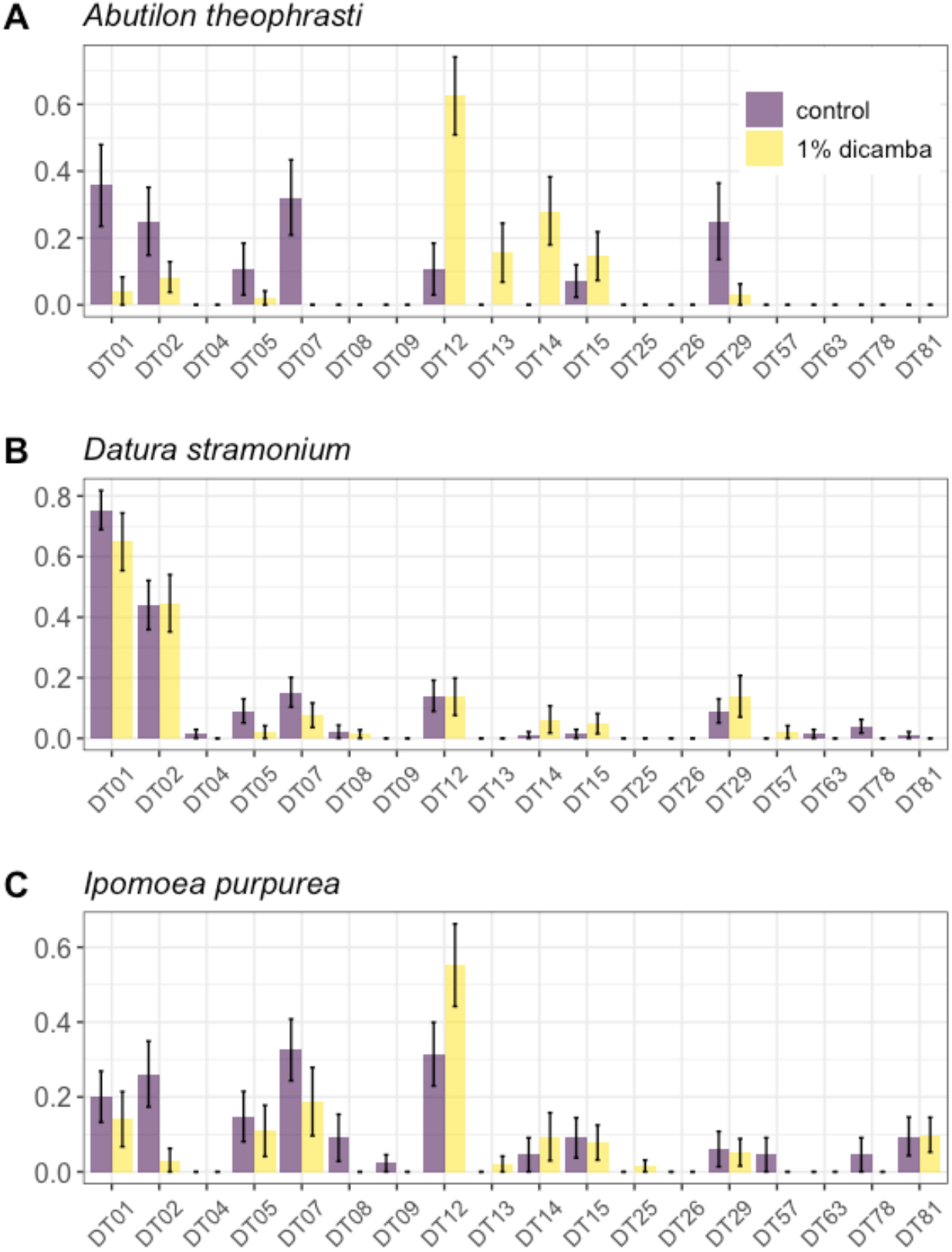
The proportion of leaves exhibiting chewing herbivory types, characterized according to the *Guide to Insect (and Other) Damage Types on Compressed Plant Fossils* (vers 3, Labandeira, et al., 2007), across replicate plants of A) *Abutilon theophrasti* (velvetleaf), B) *Datura stramonium* (jimsonweed), and C) *Ipomoea purpurea* (common morning glory) growing in field conditions and treated with 1% the field dose of the herbicide dicamba (5.6 g ai/ha) or a non-dicamba control. The type of herbivory was recorded on three or four leaves of each species, and the average proportion of each herbivory type (± standard error) across plants of each species is shown. The specific categories DT01 through DT09 are different types of hole feeding, as are DT78, DT57, DT63. DT12, DT13, DT14, DT15, DT26, and DT81 capture different types of margin feeding, and DT25 and DT29 capture surface feeding.

We further characterized the type of herbivory using the *Guide to Insect (and Other) Damage Types on Compressed Plant Fossils* (vers 3, Labandeira, et al., 2007). For *A. theophrasti*, the increase in margin feeding in the presence of dicamba appeared to be largely due to herbivores causing circular and shallow damage of the leaf margin (DT12), removal of the leaf apex (DT13), excision of the leaf to the primary vein (DT14), and excisions that are deep toward the primary vein (DT15). The increase in margin feeding across *I. purpurea* leaves appeared to be primarily caused by herbivores that left a pattern of circular and shallow damage to the leaf margin (DT12).

## DISCUSSION

In this work we found an increase in chewing herbivory damage when plants were exposed to dicamba drift, as well as changes in the type of herbivory upon exposure. Notably, we found differences between species in the amount of leaf-chewing herbivory: while *Abutilon theophrasti* and *Datura stramonium* showed more total leaf-chewing herbivory when exposed to dicamba, as compared to control plants, *Ipomoea purpurea* showed no difference in the overall amount of chewing herbivory between drift and control conditions. We also found that the type of herbivory was significantly altered in drift. *A. theophrasti* and *I. purpurea* both exhibited declines in hole feeding but increases in margin feeding, whereas *D. stramonium* showed no such changes. Additionally, surface feeding was reduced in *A. theophrasti* in the presence of dicamba, but there was no perceptible change in surface feeding according to treatment in the other two species. Thus, our expectation that normal patterns of herbivory may be disrupted in the presence of dicamba drift was met. While there were some commonalities across species, in general, patterns of chewing herbivory were different between the three weed species when exposed to dicamba.

For example, the proportion of herbivory increased in two of the three species in the presence of dicamba (*Abutilon theophrasti* and *Datura stramonium*) compared to control plants whereas the type of herbivory was altered in two species following dicamba exposure (*A. theophrasti* and *Ipomoea purpurea*). Only *A. theophrasti* exhibited both increased herbivory and a change in the type of chewing herbivory when exposed to the herbicide. Our results thus show heterogeneity in herbivory between species given dicamba exposure, and importantly, demonstrate that conclusions drawn from one weed cannot be used to generalize how dicamba drift may influence herbivory in other weeds. Further, a separate experiment using *A. theophrasti* found a nonsignificant increase in the proportion of chewing herbivory when plants were exposed to dicamba drift, again in a common garden field setting (Johnson and Baucom, 2022), potentially suggesting that results may be heterogeneous between experiments, whether due to changes in herbivore abundance or some other, unexamined environmental factor. While not chewing herbivory, this previous work also found that *A. theophrasti* exposed to dicamba drift exhibited higher abundance of whitefly larvae – a result that was replicated between two field seasons (Johnson and Baucom, 2022).

Given findings from this study and from our previous work (Johnson and Baucom, 2022), it is tempting to conclude that herbivory defense responses in *Abutilon theophrasti* may be reduced or modified when exposed to dicamba. However, we cannot determine if the changes we uncovered in the proportion or type of herbivory are due to plant- or insect-specific responses to dicamba drift. We do not know, for example, if exposure to dicamba leads to an alteration or reduction of plant defenses, allowing for greater rates of herbivory, or if dicamba reduces/alters nutrients in the plants, which may impact herbivore feeding preferences. Furthermore, the higher rates of chewing herbivory found in *A. theophrasti* and *Datura stramonium* could also be due to community changes caused by dicamba drift – perhaps predatory or parasitic insects were directly affected by dicamba exposure, such that populations of chewing herbivores were not regulated in dicamba exposed areas as they were in control plots. Although there are few studies of herbivory rates in dicamba exposed plants for comparison, examination of plant-insect interactions in plants exposed to another synthetic auxin herbicide, 2,4-D, suggest that any of the above hypotheses are possible. For example, low doses of 2,4-D on rice and corn plants led to increased growth of lepidopteran larvae (Ishii and Hirano, 1963; Oka and Pimentel, 1976) – suggesting nutrients may be altered in exposed plants – and greater population sizes of the sugarcane borer were found in 2,4-D treated sugarcane due to higher rates of oviposition and lower rates of parasitism (Ingram et al., 1947), in support of the idea that community-wide changes following herbicide exposure may occur.

The changes in the type of herbivory that we uncovered in *Ipomoea purpurea* and *Abutilon theophrasti* – *i.e*., more margin-compared to hole feeding in the presence of dicamba – could likewise be caused by dicamba-induced alterations of specific plant defenses, or changes in plant nutrients, that made dicamba exposed plants less attractive to hole feeding herbivores. Although we did not pair our quantification of herbivory with surveys of herbivore abundance across treatments, we observed a greater abundance of Japanese beetles on *A. theophrasti* control plants, and it appeared the Japanese beetles caused more hole-than margin feeding on the controls. We note, however, that Japanese beetles can cause both margin and hole feeding, and that multi-damaging capabilities are found across many herbivores (Clissold, 2007), meaning that identification of particular insect species from leaf chewing damage is not strictly possible. In further refining the type of leaf damage into 18 categories using the *Guide to Insect (and other) Damage Types on Compressed Plant Fossils* (Labandeira, et al., 2005), we found that the category of margin feeding that increased the most in both *I. purpurea* and *A. theophrasti* was DT12, ‘circular and shallow damage to the leaf margin,’ potentially suggesting that a generalist herbivore (or herbivores) shared between both weeds was affected in some way in the dicamba exposed plots. Interestingly, this specific type of margin feeding (DT12) was largely responsible for the increased margin feeding in *I. purpurea* whereas three other categories of margin feeding, in addition to DT12, were greater in *A. theophrasti* (DT13, DT14, and DT15) potentially suggesting that the latter weed was made more palatable to other herbivore species. An important next step will be to examine herbivore abundance along with patterns of chewing herbivory to understand how margin feeding might have increased on dicamba exposed plants.

Furthermore, the changes in type of herbivory that we uncovered could be due to insect life cycle differences between control and dicamba exposed plants, since changes in mouthparts due to insect life stage are known to produce particular patterns of foliar tissue damage (Hochuli, 2001).

Differences in life cycles of herbivores between treatment plots could be caused by herbivores being directly exposed to dicamba – for example, when exposed to the synthetic auxin 2,4-D, coccinellid larvae showed both delayed development and higher mortality (Adams, 1960; Michaud and Vargas, 2010). Alternatively, the herbivory type differences we uncovered could be due to indirect effects on insect development stemming from direct effects of plant exposure to dicamba. More specifically, dicamba exposure may alter the plant in some way that leads to less-than-optimal resources for herbivore growth and life cycle completion. In support of this idea, *Vanessa cardui* larvae (painted lady caterpillars) raised on dicamba exposed thistle plants were smaller compared to larvae raised on non-exposed plants (Bohnenblust et al., 2013), and similarly, when raised on 2,4-D treated plants, adult emergence of the weevil *Trichosirocalus horridus* was reduced (Stoyer and Kok, 1989). Regardless of the mechanism underlying our results, these findings show the potential of dicamba drift to alter insect herbivore community structure which may have downstream consequences, especially if dicamba exposure leads to insect growth and fitness effects that then cascade up food webs.

## CONCLUSION

Overall, our results align with work that shows differential impacts of dicamba drift across insect species (Bohnenblust et al., 2013; Egan et al., 2014b) in that we identified heterogeneous effects of dicamba drift on different crop weeds. Herbicide-induced modifications to herbivore diet and the amount of feeding may have lasting effects on ecosystems, since plant-herbivore dynamics underlie complex ecological webs that can facilitate cascading interactions (Agrawal et al., 2006; Schmitz, 2008). For instance, herbivory determines decomposition rates and nutrient cycling (Pastor and Cohen, 1997), patterns of plant assemblage (Huntly, 1991), and the frequency and growth rates of higher trophic levels (Power, 1992; Scherber et al., 2010), all of which can reciprocally influence herbivore community structure (Koricheva et al., 2000; Becerra, 2007). The results we present here demonstrate how herbicide drift can induce shifts in plant-herbivore interactions, but also highlight the need for comparative studies on communities exposed to drift in agroecosystems as well as studies that examine the mechanism behind the altered interactions.

## ACKNOWLEDGMENTS

The authors thank Michael Palmer and Jeremy Moghtader and the Matthaei Botanical Gardens and Nichols Arboretum (MBGNA) for assistance with field work. We also thank Eliot Jackson for planting and field maintenance. This work was supported by NSF DEB (1834496) and USDA (2017-09529/1016564) grants to RSB.

## AUTHOR CONTRIBUTIONS

RSB designed the experiment, performed analyses and wrote the manuscript. GZ and AS recorded herbivory on experimental leaves and helped with manuscript preparation. NJ assisted with the field work and wrote the manuscript.

## DATA AVAILABILITY STATEMENT

Data will be made available on Dryad upon acceptance.

## LITERATURE CITED

Adams, J. B. 1960. Effects of spraying 2,4-D amine on coccinellid larvae. Canadian Journal of Zoology 38: 285–288.

Agrawal, A. A. 2011. Current trends in the evolutionary ecology of plant defence. Functional Ecology 25: 420–432.

Agrawal, A. A., J. A. Lau, and P. A. Hambäck. 2006. Community heterogeneity and the evolution of interactions between plants and insect herbivores. The Quarterly Review of Biology 81: 349–376.

Baker, N. T. ‘Estimated Annual Agricultural Pesticide.’ 1992–2019 United States Geological Survey. Website https://water.usgs.gov/nawqa/pnsp/usage/maps/show_map.php?year=2019&map=DICAMBA&hilo=L&disp [accessed 17 August 2022].

Bates, D., M. Maechler, and B. Bolker. 2011. lme4: linear mixed-effects models using S4 classes.

Baucom, R. S. 2019. Evolutionary and ecological insights from herbicide-resistant weeds: what have we learned about plant adaptation, and what is left to uncover? The New Phytologist.

Becerra, J. X. 2007. The impact of herbivore-plant coevolution on plant community structure. Proceedings of the National Academy of Sciences of the United States of America 104: 7483–7488.

Bohnenblust, E., J. F. Egan, D. Mortensen, and J. Tooker. 2013. Direct and indirect effects of the synthetic-auxin herbicide dicamba on two lepidopteran species. Environmental Entomology 42: 586–594.

Carvalho, M. R., P. Wilf, H. Barrios, D. M. Windsor, E. D. Currano, C. C. Labandeira, and C. A. Jaramillo. 2014. Insect leaf-chewing damage tracks herbivore richness in modern and ancient forests. PloS One 9: e94950.

Cessna, Wolf, and Stephenson. 2005. Pesticide movement to field margins: routes, impacts and mitigation. Insect and Disease Management 1: 69–112.

Clissold, F. J. 2007. The Biomechanics of Chewing and Plant Fracture: Mechanisms and Implications. In J. Casas, and S. J. Simpson [eds.], Advances in Insect Physiology, 317–372. Academic Press.

Cyr, H., and M. L. Face. 1993. Magnitude and patterns of herbivory in aquatic and terrestrial ecosystems. Nature 361: 148–150.

Egan, J. F., K. M. Barlow, and D. A. Mortensen. 2014a. A Meta-Analysis on the Effects of 2,4-D and Dicamba Drift on Soybean and Cotton. Weed Science 62: 193–206.

Egan, J. F., E. Bohnenblust, S. Goslee, D. Mortensen, and J. Tooker. 2014b. Herbicide drift can affect plant and arthropod communities. Agriculture, Ecosystems & Environment 185: 77–87.

EPA. 2021. Status of Over-the-Top Dicamba: Summary of 2021 Usage, Incidents and Consequences of Off-Target Movement, and Impacts of Stakeholder-Suggested Mitigations (DP# 464173: PC Code 128931). American Library Association.

Forister, M. L., V. Novotny, A. K. Panorska, L. Baje, Y. Basset, P. T. Butterill, L. Cizek, et al. 2015. The global distribution of diet breadth in insect herbivores. Proceedings of the National Academy of Sciences of the United States of America 112: 442–447.

Fuchs, B., K. Saikkonen, and M. Helander. 2020. Glyphosate-Modulated Biosynthesis Driving Plant Defense and Species Interactions. Trends in Plant Science.

Futuyma, D. J., and A. A. Agrawal. 2009. Macroevolution and the biological diversity of plants and herbivores. Proceedings of the National Academy of Sciences of the United States of America 106: 18054–18061.

Grossmann, K. 2010. Auxin herbicides: current status of mechanism and mode of action. Pest Management Science 66: 113–120.

Gutiérrez, Y., D. Ott, and C. Scherber. 2020. Direct and indirect effects of plant diversity and phenoxy herbicide application on the development and reproduction of a polyphagous herbivore. Scientific Reports 10: 7300.

Hochuli, D. F. 2001. Insect herbivory and ontogeny: How do growth and development influence feeding behaviour, morphology and host use? Austral Ecology 26: 563–570.

Huntly, N. 1991. Herbivores and the Dynamics of Communities and Ecosystems. Annual Review of Ecology and Systematics 22: 477–503.

Ingram, J. W., E. K. Bynum, L. J. Charpentier. 1947. Effect of 2, 4-D on sugarcane borer. Journal of Economic Entomology 40: 745–746.

Ishii, S., and C. Hirano. 1963. Growth responses of larvae of the rice stem borer to rice plants treated with 2, 4-D. Entomologia Experimentalis et Applicata.

Johnson, N. M., and R. S. Baucom. 2022. Dicamba drift alters plant-herbivore interactions at the agro-ecological interface. bioRxiv: 2021.07.13.452219.

Jones, G. T., J. K. Norsworthy, T. Barber, E. Gbur, and G. R. Kruger. 2019. Off-target Movement of DGA and BAPMA Dicamba to Sensitive Soybean. Weed Technology: A Journal of the Weed Science Society of America 33: 51–65.

Koricheva, J., C. P. H. Mulder, B. Schmid, J. Joshi, and K. Huss-Danell. 2000. Numerical responses of different trophic groups of invertebrates to manipulations of plant diversity in grasslands. Oecologia 125: 271–282.

Labandeira, C.C., Wilf, P., Johnson, K.R., and Marsh, F. 2007. Guide to insect (and other) damage types on compressed plant fossils. Version 3.0. Smithsonian Institution, Washington.

Liu, J., and X.-J. Wang. 2006. An integrative analysis of the effects of auxin on jasmonic acid biosynthesis in Arabidopsis thaliana. Journal of Integrative Plant Biology 48: 99–103.

Massad, T. J., and L. A. Dyer. 2010. A meta-analysis of the effects of global environmental change on plant-herbivore interactions. Arthropod-Plant Interactions 4: 181–188.

Michaud, J. P., and G. Vargas. 2010. Relative toxicity of three wheat herbicides to two species of Coccinellidae. Insect Science.

Miles, L. S., S. T. Breitbart, H. H. Wagner, and M. T. J. Johnson. 2019. Urbanization shapes the ecology and evolution of plant-arthropod herbivore interactions. Frontiers in Ecology and Evolution 7.

Muday, G. K., A. Rahman, and B. M. Binder. 2012. Auxin and ethylene: collaborators or competitors? Trends in Plant Science 17: 181–195.

Oka, I. N., and D. Pimentel. 1976. Herbicide (2,4-D) Increases Insect and Pathogen Pests on Corn. Science 193: 239–240.

Owen, M. D. K. 2008. Weed species shifts in glyphosate-resistant crops. Pest Management Science 64: 377–387.

Pastor, J., and Y. Cohen. 1997. Herbivores, the Functional Diversity of Plants Species, and the Cycling of Nutrients in Ecosystems. Theoretical Population Biology 51: 165–179.

Peña-Cortés, H., J. Fisahn, and L. Willmitzer. 1995. Signals involved in wound-induced proteinase inhibitor II gene expression in tomato and potato plants. Proceedings of the National Academy of Sciences of the United States of America 92: 4106–4113.

Peterson, R. Ryan, and A. Peterson. 2021. Finding Optimal Normalizing Transformations via bestNormalize. The R Journal 13: 310.

Power, M. E. 1992. Top-Down and Bottom-Up Forces in Food Webs: Do Plants Have Primacy. Ecology 73: 733–746.

Ramankutty, N., Z. Mehrabi, K. Waha, L. Jarvis, C. Kremen, M. Herrero, and L. H. Rieseberg. 2018. Trends in Global Agricultural Land Use: Implications for Environmental Health and Food Security. Annual Review of Plant Biology 69: 789–815.

RStudio Team. 2020. RStudio: Integrated Development for R. RStudio, PBC, Boston, MA.

Satterthwaite, F. E. 1946. An approximate distribution of estimates of variance components. Biometrics 2: 110–114.

Scherber, C., N. Eisenhauer, W. W. Weisser, B. Schmid, W. Voigt, M. Fischer, E.-D. Schulze, et al. 2010. Bottom-up effects of plant diversity on multitrophic interactions in a biodiversity experiment. Nature 468: 553–556.

Schindelin, J., I. Arganda-Carreras, E. Frise, V. Kaynig, M. Longair, T. Pietzsch, S. Preibisch, et al. 2012. Fiji: an open-source platform for biological-image analysis. Nature Methods 9: 676–682.

Schmitz, O. J. 2008. Herbivory from Individuals to Ecosystems. Annual Review of Ecology, Evolution, and Systematics 39: 133–152.

Sharma, A., P. Jha, and G. V. P. Reddy. 2018. Multidimensional relationships of herbicides with insect-crop food webs. The Science of the Total Environment 643: 1522–1532.

Stoyer, T. L., and L. T. Kok. 1989. Oviposition by Trichosirocalus horridus (Coleoptera: Curculionidae), a Biological Control Agent for Carduus Thistles, on Plants Treated with Low Dosages of 2,4-Dichlorophenoxyacetic Acid. Environmental Entomology 18: 715–718.

Teale, W. D., I. A. Paponov, and K. Palme. 2006. Auxin in action: signaling, transport and the control of plant growth and development. Nature reviews. Molecular Cell Biology 7: 847–859.

Team, R. C. 2016. R: A language and environment for statistical computing. R Foundation for Statistical Computing, Vienna, Austria. 2013.

Tilman, D., P. B. Reich, and F. Isbell. 2012. Biodiversity impacts ecosystem productivity as much as resources, disturbance, or herbivory. Proceedings of the National Academy of Sciences of the United States of America 109: 10394–10397.

Turcotte, M. M., H. Araki, D. S. Karp, K. Poveda, and S. R. Whitehead. 2017. The eco-evolutionary impacts of domestication and agricultural practices on wild species. Philosophical Transactions of the Royal Society of London. Series B, Biological Sciences 372.

Turcotte, M. M., T. Jonathan Davies, C. J. M. Thomsen, and M. T. J. Johnson. 2014. Macroecological and macroevolutionary patterns of leaf herbivory across vascular plants. Proceedings of the Royal Society B: Biological Sciences 281: 20140555.

Wechsler, Smith, and McFadden. 2019. The use of genetically engineered dicamba-tolerant soybean seeds has increased quickly, benefiting adopters but damaging crops in some fields. Amber Waves: the economics of food, farming, natural resources and rural America. https://www.ers.usda.gov/amber-waves/2019/october/the-use-of-genetically-engineered-dicamba-tolerant-soybean-seeds-has-increased-quickly-benefiting-adopters-but-damaging-crops-in-some-fields/

Wickham, H. 2016. Data Analysis. In H. Wickham [ed.], ggplot2: Elegant Graphics for Data Analysis, 189–201. Springer International Publishing, Cham.

Xu, P., P.-X. Zhao, X.-T. Cai, J.-L. Mao, Z.-Q. Miao, and C.-B. Xiang. 2020. Integration of Jasmonic Acid and Ethylene Into Auxin Signaling in Root Development. Frontiers in Plant Science 11: 271.

